# Phylogeny, transposable element and sex chromosome evolution of the basal lineage of birds

**DOI:** 10.1101/750109

**Authors:** Zongji Wang, Jilin Zhang, Xiaoman Xu, Christopher Witt, Yuan Deng, Guangji Chen, Guanliang Meng, Shaohong Feng, Tamas Szekely, Guojie Zhang, Qi Zhou

**Author notes:** These authors contributed equally to the work (Zongji Wang, Jilin Zhang). Correspondence should be addressed to Q. Z. or G. Z.

## Abstract

Sex chromosomes of mammals and most birds are heteromorphic, while those of many paleognaths (ratites and tinamous) are inexplicably homomorphic. To dissect the mechanisms underlying the different tempo of sex chromosome evolution, we produced high-quality genomes of 12 paleognathous species, and reconstructed their phylogeny based on alignments of the non-coding sequences extending to nearly 40% of the genome. Our phylogenomic tree grouped the South American rheas and tinamous together, and supported the independent evolution of gigantism and loss of flight among ratites. The small-bodied tinamous have much higher rates of genome-wide substitutions and transposon turnovers. Yet majorities of both have retained exceptionally long recombining regions occupying over half of the entire sex chromosome, with the rest sex-linked regions diverging from each other at a much lower rate relative to neognathous birds. Each species exhibits a punctuated sequence divergence pattern between sex chromosomes termed ‘evolutionary strata’, because of stepwise suppression of recombination. We concluded that all paleognaths share one evolutionary stratum with all other birds, and convergently formed between one to three strata after their rapid speciation. Contrary to the classic notion, we provided clear evidence that the youngest stratum of some tinamous formed without chromosomal inversion. Intriguingly, some of the encompassing W-linked genes have upregulated their expression levels in ovary, probably due to the female-specific selection. We proposed here that the unique male-only parental care system of paleognaths has reduced the intensity of sexual selection, and contributed to these species’ low rates of sex chromosome evolution. We also provided novel insights into the evolution of W-linked genes at their early stages.

## Introduction

Palaeognathae comprises less than 1% of all avian species, but has been intriguing biologists for over a century with an elusive evolutionary history and many extraordinary features that are informative for the origin and evolution of birds in general. As the basal branch to all the rest living birds (Neognathae), it is traditionally divided into two groups: the usually gigantic (except for kiwis) flightless ratites and the small pheasant-like volant tinamous ^1,2^. Most Palaeognathae fossils were discovered in the Northern Hemisphere ^3^. Thus, the remarkable distribution of extant ratites (e.g., kiwi and rhea) in the Southern Hemisphere was previously taken as a textbook demonstration of vicariance biogeography. It was hypothesized that the divergence from the single flightless common ancestor of ratites was driven by the breakup of Gondwana and the following continental drift ^4^. Such a ratite monophyly was mainly based on their morphological and behavioral traits, and has been challenged by molecular phylogenetic analyses using mitochondrial and nuclear gene sequences. Mounting evidence supporting the ratite polyphyly recently emerged from the phylogenomic studies using the recovered ancient DNAs from extinct elephant birds and moa ^5–7^. The Madagascan elephant birds were grouped together with the New Zealand kiwi, and the South American tinamous with the New Zealand moa, whose living landmasses were never connected in Gondwana. These results are not expected by the vicariant divergence model, which predicts for example, the elephant birds to be grouped with ostriches in Africa. Much progress has been made in resolving the Palaeognathae phylogeny. Different studies now consistently placed the ostrich basal to all other paleognaths, and the emu, cassowary, and kiwis as one group, but have reported various phylogenetic positions for rheas due to the ancestral incomplete lineage sorting (ILS) ^1,6–8^ (**Supplementary Table S1**). These studies also consistently placed the flighted tinamous within rather than completely separated from all ratites, indicating flight loss and gigantism have convergently evolved multiple times during the ratite divergence through flight dispersal.

The evolution of associated traits of flight loss and gigantism has facilitated ratites’ adaption for cursoriality, and had profound impacts on their genome evolution. Compared to the genomes of other smaller and volant birds, the ostrich genome is characterized with much lower rates of substitutions ^9^ and insertions/deletions, transposon removal ^10^, and very few intrachromosomal rearrangements even when being compared with the distantly-related chicken^11^. This parallels the similar comparisons between elephants or whales vs. other mammals ^10,12–14^, and can be explained by the fact that species with a larger body size tend to have a longer generation time and a lower metabolic rate, hence a reduced mutation rate ^12,15^. Much fewer lineage-specific changes and the early-branching feature relative to the majority of birds make the large-bodied ratites a great model for reconstructing the trajectory of evolution from the avian ancestor. It may explain why the sex chromosomes of many ratites are arrested in their ancestral states, and are still evolving largely like a pair of autosomes since their origination over 100 million years (MY) ago ^13,16,17^. In contrary to the XY sex system of mammals, all birds share one pair of homologous female heterogametic sex chromosomes (male ZZ, female ZW). We have previously reported that most Neognathae species have undergone four times of recombination suppression (RS) between sex chromosomes, with each influenced region forming a distinct pattern (termed ‘evolutionary strata’ ^18^) of Z/W pairwise sequence divergence level from those of nearby regions^16^. As a result, over 90% sex-linked regions of most birds^16,19^, like those of all mammals^20^ and many other vertebrates, are highly differentiated from each other except for the chromosome ends as the pairing pseudoautosomal regions (PAR). Emu and ostrich present a striking exception, and have over two-thirds of the sex-linked region as PAR, after only two times of RS^16,21^. A lower mutation rate, at both the genome-wide level due to their large body size, and on the female-specific W chromosome due to the ‘male-driven evolution’ effect ^18,22^ may account for the lower rate of sex chromosome divergence of ratites. Alternatively, it can be associated with the unique parental care system of paleognaths, that males exclusively or predominantly care for the offspring while females desert the eggs, in contrast to other 95% of bird species with biparental or female-only care^23^. Such a male-only care system has probably originated in birds’ dinosaur ancestors ^24^ before the origination of avian sex chromosomes, and is associated with the large clutch size, i.e., female premating investment. It may balance the post-mating sexual conflict over care and result in a lower intensity of premating sexual selection targeting males, indicated by paleognaths’ much lower degree of sexual dimorphism compared to most other bird species ^25,26^ (**Supplementary Table S2**). Overall, reduced sexual conflict and sexual selection may account for the unique sex chromosome pattern of ratites^27,28^.

These hypotheses concern the major forces driving the different tempo of sex chromosome evolution. To clarify the effects of mutation vs. sexual selection, and also to provide deep insights into the genome evolution of Palaeognathae in general, we here produced female genomes of 9 tinamou species and 3 ratite species, and analyzed a total of 15 Palaeognathae genomes. We first placed rhea next to the tinamou clade with high confidence level, based on a phylogenomic tree reconstructed from non-coding sequence alignments extending to nearly 40% of the entire genome length per species. Then we characterized most tinamou species, despite a much longer branch length and faster turnovers of transposons than ratites, show a similar pattern of sex chromosome evolution comparing to ratites. This suggests that the reversed sex role in parental-care of Palaeognathae probably play an important role besides the mutation rates in shaping their patterns of sex chromosome evolution.

## Results

### Phylogenomic analyses group rhea with tinamous

Previous works have placed rheas as the sister branch to all paleognaths except for ostrich ^6,29^, or to the clade of emus, cassowaries and kiwis ^7,8^, or to the clade of tinamous ^1,8^, many of which have a low bootstrapping score at the rhea node (**Supplementary Table S1**). To resolve this last contentious node in the Palaeognathae phylogeny, we generated high-quality genome assemblies and annotations for 12 paleognathous species, from between 48 and 106 fold of shotgun sequencing per species (**Supplementary Table S3**). We specifically chose female samples of all species for sequencing to include the genomic information of W chromosomes, and densely sampled tinamou species (9 species) to form a comprehensive comparison to ratites, as well as for tinamous’ intriguingly diverse sex chromosome composition^21,30^ and unknown genome evolution patterns. Two thirds of the sequenced species have a draft genome with a scaffold N50 length longer than 1Mb, and for the assembled regions, all genomes have a gap content lower than 5%, comprising on average 81.52% of the estimated genome size. For each genome, we annotated from 13,150 to 16,438 protein-coding genes, and about 5% of the genome as repetitive elements, with an average BUSCO score reaching 92.6 (**Supplementary Table S4-5**). We have also assembled the complete mitochondrial genomes, and annotated a complete set of 13 mitochondrial genes of all the studied species.

It is known that the topology of phylogenetic tree is sensitive to the amount and the type of data, the number of sampled species, as well as the method of phylogenetic inference ^9,31,32^. We therefore generated phylogenomic trees based on the concatenated alignments of whole-genome non-coding sequences (NC, **Figure 1A**), of entire mitochondrial genomes (MT, **Figure. 1B**), and of fourfold degenerate sites from single-copy orthologous genes (4D, **Figure 1C**), with ostrich fixed as the sister branch to all the other paleognaths. After removing alignment ambiguities, these datasets comprise per species 393,165,276 bp noncoding sequences (~37.56 % of the nuclear genome), 15,873bp or 92.57% of the mitochondrial genome, and 588,067 bp from 4,387 orthologous genes or 30.78% of the entire gene repertoire. They are expectedly larger than the previous whole-genome alignments involving 48 avian species that are more divergent from each other ^9^, and represent the largest phylogenomic dataset of Palaeognathae known to date (**Supplementary Table S1**). Given the species trees reconstructed by coalescence (CL) method or presence/absence of transposons (TE) have been recently published ^7^, for comparison we produced here the maximum likelihood (ML) trees for the MT and NC datasets, and the Bayesian tree for the 4D dataset. All resulting phylogenies have reached 100% full bootstrapping support for all the nodes of the studied species. Topologies of NC-tree and MT-tree are consistent with each other and have placed the greater rhea as the sister species to all tinamou species, with both taxa now living in South America. This is different from the 4D Bayesian tree, and a previous Palaeognathae ML tree based on a much smaller dataset of 27.9Mb intron alignments, which placed rheas as a sister lineage to all non-ostrich paleognaths ^7^. It is also different from the published CL tree which grouped rheas with the clade of emu/cassowary/kiwis ^7^. The difference between the CL- and NT-trees can be caused by the different data types and methods of phylogenetic inferences^31^. While the difference between 4D- and NT-trees of this work can reflect the influence on the gene sequences from convergent evolution of life history traits, and the dataset of 4D-tree is rather small. So we used the NC-tree topology as the guide tree for all subsequent analyses in this work. Our estimation of species divergence time is generally consistent, with some differences in the tinamou lineage compared with previous results (**Supplementary Fig. S1**)^5,6^, because of much more dense samplings of tinamou species in this work. Particularly, the estimated divergence time (about 67.9 MY) of rhea from its sister group is around the Cretaceous– Paleogene (K-Pg) mass extinction 66 MY ago ^9^, when high levels of ILS also occurred during speciation of Neoaves. Overall, the branch lengths of ratites are much shorter than those of tinamous (**Figure 1A**, *P* =1e-07, relative rate test), indicating a much lower genomic substitution rate among ratites due to the evolution of gigantism.

**Figure 1.**
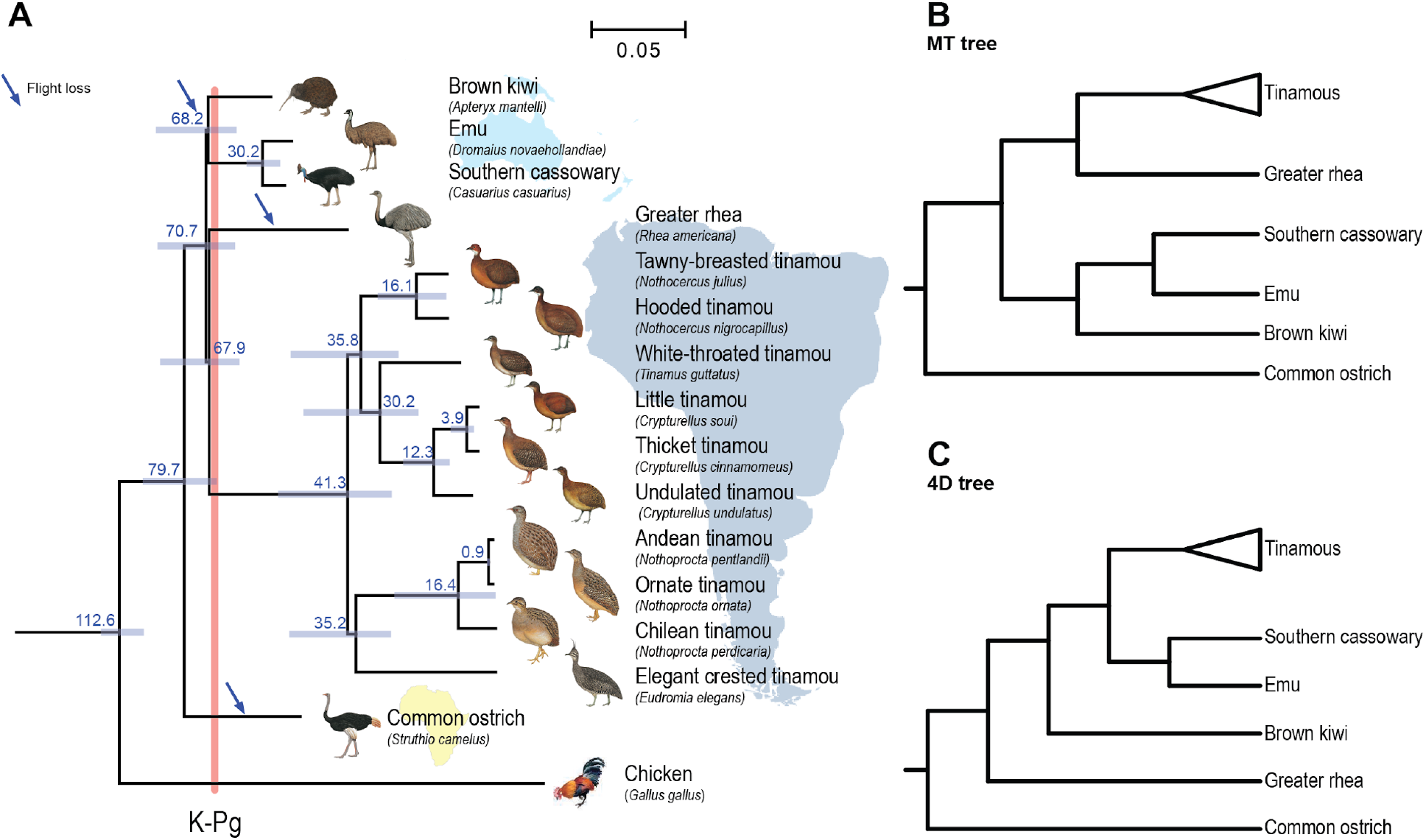
Phylogenomic tree of Palaeognathae. (A) Maximum likelihood tree reconstructed from genome-wide alignments of non-coding sequences. Branch lengths indicating the genome substitution rates were estimated by ExaML. All phylogenetic nodes have full bootstrap support, and were labelled with estimated species divergence times (in million years), and 95% highest posterior densities (in blue horizontal bars). Based on the tree, all extant paleognaths are grouped by their living landmass shown in the background, and extant ratites probably have had at least three times of independent flight loss marked by blue arrows. The divergence time between rheas and tinamous is very close to the Cretaceous-Paleogene (K-Pg, in red bar) mass extinction about 66 million years ago. All bird illustrations were ordered from https://www.hbw.com/ ^90^. (B) Bayesian tree constructed with fourfold degenerate sites from protein-coding genes. (C) Maximum likelihood tree reconstructed from the concatenation of mitochondrial genomes.

### Temporal evolution of transposons and population sizes of paleognaths

It has been proposed that the metabolic requirement for powered flight is linked to a reduced genome size ^10,33^. Indeed, with a similar sequencing coverage, the assembled genome sizes of ratites are on average 27.72% larger (*P*=0.0012, Wilcoxon test) than those of flighted tinamous (**Supplementary Table S4**). Besides the reported different rates of genomic deletions between the two groups of species, the contribution of transposable element (TE) turnovers to such variable genome sizes of paleognaths remains to be elucidated, as previous analysis included only two species of ostrich and white-throated tinamou^10^. Here we found that paleognaths generally have a similar level of genome-wide repeats, ranging from 4.89% of thicket tinamou to 5.54% of ostrich, which is largely attributed to their similar long interspersed nuclear element (LINE) content. Ratites have significantly (*P*<0.0007, Wilcoxon test) more DNA transposons (on average 0.55 fold higher) and short interspersed nuclear elements (SINE, on average 1.25 fold higher) than those of tinamous. They together have contributed to on average additional 13.98 Mb sequences to the ratite genomes compared to those of tinamous (**Supplementary Table S6**). When inspecting the sequence divergence patterns of DNA transposons comparing to their consensus sequences (**Supplementary Fig. S2**), we have not found any recent expansions in ratites, whose divergence level is expected to be low (**Figure 2A**). Thus the higher DNA transposon content of ratites is more likely to be caused by the higher rate of DNA transposon removal in tinamous.

**Figure 2.**
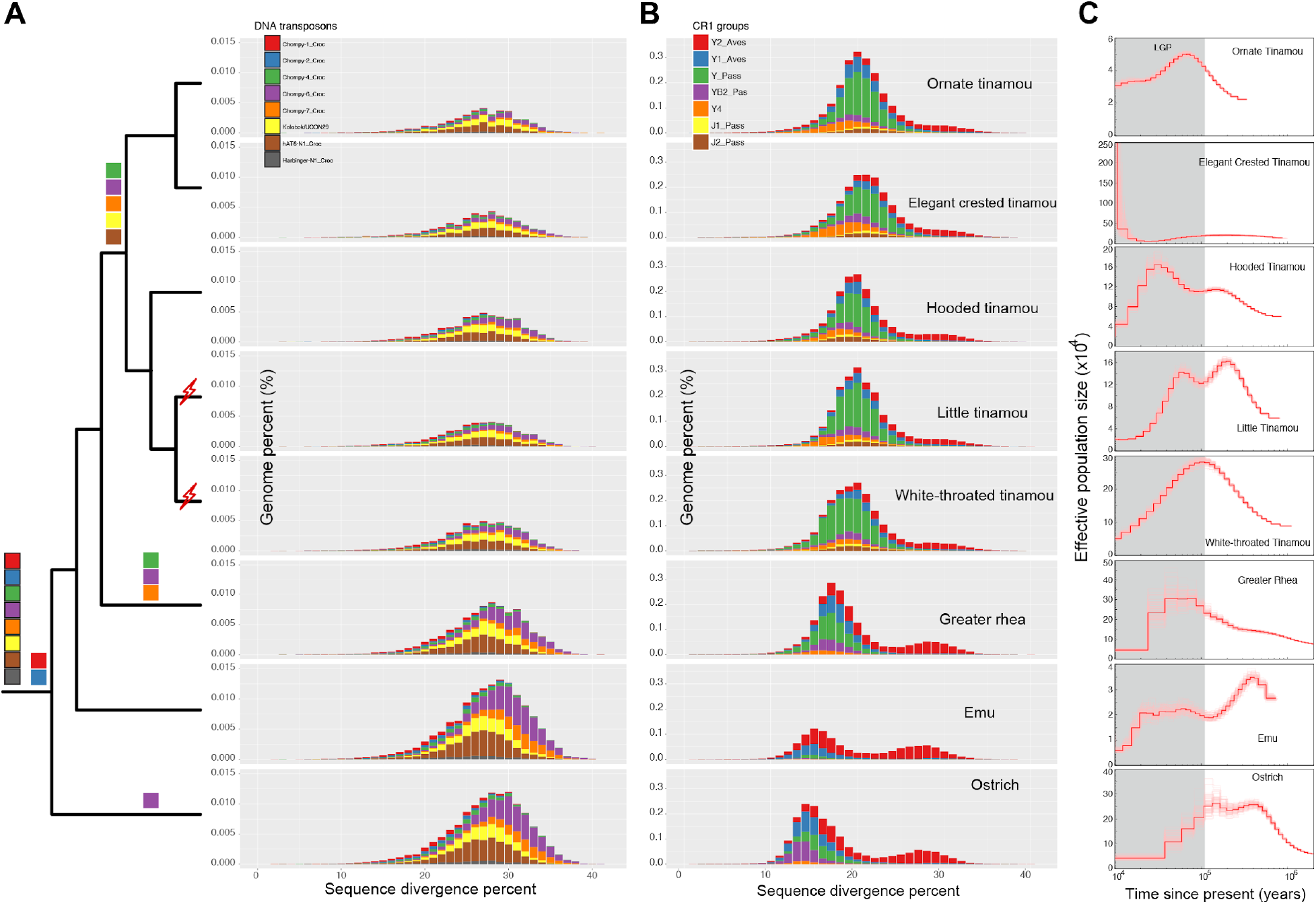
Temporal evolution of transposons and population sizes of paleognaths. (A, B) We labelled the inferred bursts of certain subfamilies of DNA transposons (in black-framed color squares) and CR1 LINEs (in squares without black frames) at the corresponding phylogenetic nodes. The histograms show the distributions of sequence divergence between each subfamily vs. their consensus sequences. We also dated two horizontal transfer of AviRTE retroposon, one of which has been reported previously^35^, indicated here by the red lighting signs. (C) Dynamics of population size of paleognaths inferred from PSMC analyses in the past 1 million years, with variations in population size shown as the pink curves derived from 100 bootstraps. The gray shaded areas indicate the last glacial period (LGP).

In contrast, the predominant repeat type (about 3% of the genome) chicken repeat 1 (CR1) LINE elements show a similar genomic percentage among paleognaths (**Supplementary Table S6**). Different subfamilies of CR1 repeats exhibit different patterns of genomic composition and temporal activities between ratites and tinamous. Certain subfamilies, for example, CR1-Y1_Aves and -Y2_Aves were estimated to be active throughout the evolutionary history of paleognathous species (**Supplementary Fig. S3**), and probably throughout that of all birds, as they have been previously estimated to be active in chicken and zebra finch as well^34^. This is based on our chronological analyses of nested TEs, with the expectation that younger and active TEs are more likely to be nested within older and inactive TEs (‘transposition in transposition’, TinT) than the opposite scenario. While subfamilies CR1-Y_Pass, -Y4, -YB2_Pass etc. were probably active more recently, mostly throughout the tinamou lineage, as well as in some ratites (e.g., CR1-YB2_Pass in ostrich) from the TinT analyses (**Supplementary Fig. S4**). This is corroborated by their sequence divergence pattern compared to their consensus sequences (**Figure 2B**): recently active CR1 subfamilies (e.g., CR1-Y4) are specifically enriched in tinamou species and show a sequence divergence pattern concentrated at a lower divergence level than that of those (e.g., CR1-Y2_Aves) active at an earlier time point. Overall, the lower contraction rate of ancient CR1 elements in ratites, and more recent expansion of some CR1 subfamilies in tinamou together resulted in the generally similar level of LINE elements between the two groups of species. It is worth noting that we also dated two independent horizontal transfers of retroposon, AviRTE from filarial nematodes to tinamou species, which is absent in any other paleognaths. One of them was previously reported in white-throated tinamou^35^. Based on comparison of the orthologous flanking sequences of AviRTE among species, we inferred another horizontal transfer in the ancestor of little tinamou and undulated tinamou, within 30 MY (**Supplementary Fig. S5** and **Supplementary Table S7**).

The different TE content of paleognathous species can be impacted by their different effective population sizes (Ne) ^36,37^. This has been demonstrated in the endangered mammals that recent bottlenecks can fix excessive TEs in the genome by genetic drift ^38,39^. To inspect the relationship between the TE content and the population size, we reconstructed the dynamic changes of population size of all the studied species in the past 10,000 to 100,000 years using the pairwise sequentially Markovian coalescent (PSMC) approach^38^. Sister species, for example, emu and cassowary, ornate and Andean tinamous show very similar trajectories of population size changes, probably due to their overlapped or close ecological niche after their speciation (**Supplementary Fig. S6**). Similar to the reported results ^40^, most species except for Chilean and elegant-crested tinamous, show a decline in population sizes since the beginning of last glacial period (LGP) about 100,000 years ago (**Figure 2C**). The population size does not have a significant association with the overall or individual families of TE abundance of the studied paleognathous species (**Supplementary Table S8**). For example, the dramatic recent expansion of population size of elegant-crested tinamou, or contraction of that of emu, does not result in dramatically different genome-wide TE content between the two species (5.02% vs. 4.98%). However, the recent population size change may impact the frequencies of TE families within population, as reported in Drosophila^41^.

### Halted recombination suppression and diverse evolution history of palegnathous sex chromosomes

We previously determined that about two thirds of the ostrich and emu sex chromosomes have halted the progression of RS as PAR^16^. A recent study^21^ reported that two tinamou species also have a long PAR. However, the great diversity of PAR lengths among tinamous implicated by cytogenetic studies ^30^, and particularly the diversity of W-linked genes of paleognaths remain to be elucidated. To reconstruct a more complete history of sex chromosome evolution of paleognaths, we first determined the length of PAR for each studied species, where the aligned read depth is expected to show a similar level between autosomes and sex chromosomes, or between sexes. Among the 15 studied paleognaths, 9 species, particularly half of the investigated tinamous have a PAR extending at least half of the entire sex chromosome length (**Figure 3A**). This indicated that a reduced evolution rate associated with gigantism alone cannot explain the unusual pattern of Palaeognathae sex chromosomes, as tinamous have a small body size and a similarly long branch length comparable to chicken (**Figure 1A**).

**Figure 3.**
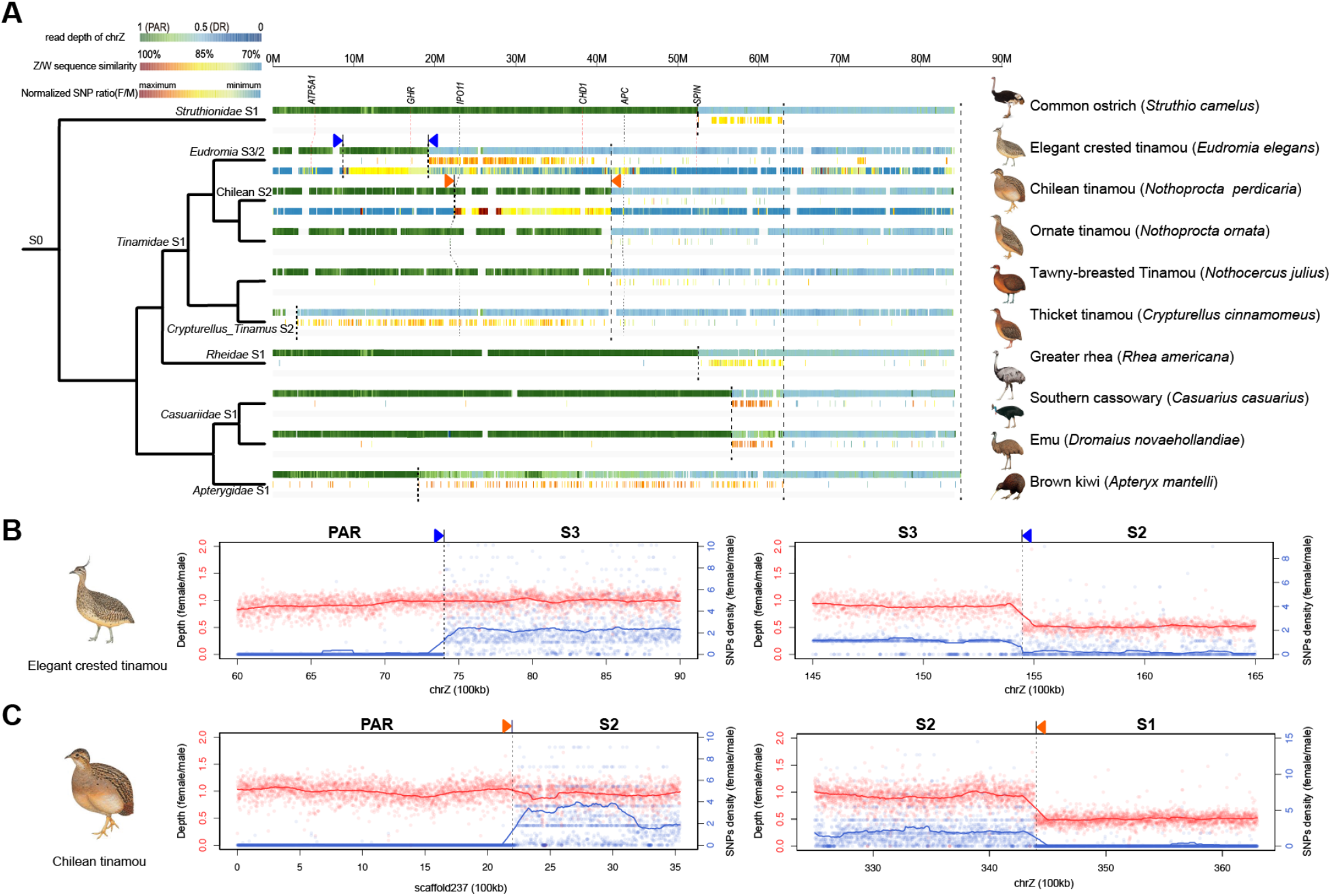
Independent evolution of evolutionary strata in Palaeognathae. (A) Each species has the first two tracks showing the color-scaled female read depths along the Z chromosome normalized against the median value of the read depths of autosomes, and the level of Z/W pairwise sequence divergence. PAR region is expected to show a normalized female read depth around 1 (in green), while the SDR whose recombination has been suppressed is expected to show a normalized read depth around 0.5 (in light blue). The faster one SDR region is evolving between sex chromosomes, the less W-linked fragments can be identified, and the more diverged in sequence that homologous Z/W regions are. For elegant-crested tinamou and Chilean tinamou, we used a third color-scaled track to show the female-to-male ratios of SNPs. An equal ratio of SNPs between sexes is expected for the PAR, while abundant female-specific heterozygotes suggest a very young evolutionary stratum, despite the influenced region show an equal read depth between sexes. The boundaries between two neighbouring evolutionary strata were inferred by the sharp shift of the Z/W divergence level or the identified W-linked fragments, or that of the female read depth. We named each evolutionary stratum according to the phylogenetic branch that it originated, and labelled them on the species tree. All bird illustrations were ordered from https://www.hbw.com/ ^90^. (B,C). Distributions of female-to-male read depth ratios (in red) and SNP ratios (in blue) around the boundaries of evolutionary strata in elegant crested tinamou (B) and Chilean tinamou (C). The positions of strata regions are marked by blue triangles in elegant crested tinamou, and by orange triangles in Chilean tinamou in (A).

In the sex-differentiated regions (SDR) where recombination has been suppressed, the density of assembled W-linked fragments, and their pairwise sequence divergence level with the homologous Z-linked regions are not uniformly distributed along the Z chromosome, forming a pattern of ‘evolutionary strata’ (**Supplementary Fig. S7**). We demarcated the strata by the sharp shift of the densities of assembled W-linked fragments, or that of the levels of Z/W divergence. We inferred a more ancestral evolutionary stratum if its boundaries are shared across multiple species, otherwise as a more recent lineage-specific stratum. An older evolutionary stratum is also expected to have less assembled W-linked fragments and a lower sequence similarity between Z/W chromosomes, as it has undergone longer time of divergent evolution between sex chromosomes. We first determined the evolutionary stratum shared by all birds (stratum 0, S0) encompassing the avian male-determining gene *DMRT1* ^16^, which emerged over 100 MY ago and is also the stratum shared by all paleognaths. After their divergence, ostrich, the ancestor of all tinamous, rhea, the ancestor of emu and cassowary, and kiwi have independently experienced a second RS event that influenced different lengths of regions and formed a stratum (S1) 23 to 75 MY ago (**Supplementary Table S9**). As we have only one species of rhea and kiwi, this S1 could have formed either in the ancestors of all rheas and all kiwis respectively, or specifically in the species that we studied here. Some tinamou species, i.e., the ancestor of *Crypturellus* genus (including thicket, little, undulated, and white-throated tinamous), elegant-crested tinamou have independently formed a S2 (**Supplementary Fig. S7**) 8 to 56 MY ago.

Intriguingly, parts of the supposed PARs of elegant-crested tinamou and Chilean tinamou exhibit a female-specific elevation of heterozygosity level, i.e., an increased divergence level between Z and W chromosomes, despite that their levels of read coverage are equal between sexes. Such regions are bordered with the two species’ S2 or S1, and the ‘true’ PARs with both the read coverage and heterozygosity levels equal between sexes (**Figure 3B-C**). They are very likely an independent S3 or S2 in the two species that have originated within 31.6 and 8 MY, respectively according to the species divergence time (**Supplementary Fig. S1, Supplementary Table S9**). It is usually assumed, and in fact has been demonstrated in human that evolutionary strata emerged through chromosomal inversions ^18,42,43^. However, previous fluorescence *in situ* hybridization (FISH) in elegant-crested tinamou showed that two genes, *ATP5A1* in the PAR, and *GHR* in the S3 have exactly the same syntenic gene order between the Z and W chromosomes ^44^. This suggests the S3 of elegant-crested tinamou formed without chromosome inversions being involved. Overall, we have reconstructed the complex history of Palaeognathae sex chromosome evolution: all the species share one RS event at S0. Because of the following independent formation of between one to three evolutionary strata after the species divergence, different species have different lengths of but generally long PARs.

### Evolution of sex-linked genes and transposable elements of Palaeognathae

We further confirmed the origination time of evolutionary strata by phylogenetic analyses of Z-or W-linked genes. We have annotated between 550 to 930 Z-linked genes and 8 to 135 W-linked genes across the studied species (**Supplementary Table S9**). The drastic differences in numbers of genes between Z and W chromosomes reflect the degeneration of W chromosomes after recombination was suppressed. If Z- and W-linked genes (also called ‘gametologs’) of certain evolutionary strata had started to diverge before the species divergence, i.e., in the case of a shared stratum across species, we expected their gametolog trees to be grouped by either Z or W gametolog sequences. Otherwise, for a lineage-specific evolutionary stratum, we expected the Z/W gametologs of the same species to be grouped together (**Supplementary Fig. S8-S13**). The ages of evolutionary strata inferred from gametolog trees are consistent with those inferred from aligning their boundaries. For example, all tinamou species share one evolutionary stratum (Tinamidae S1, named after its phylogenetic node when it originated, **Figure 3A** and **4A**), whose boundaries are aligned between tinamous by the nearby PAR or S2 sequences. Consistently, gametologs of *APC* gene from this stratum are grouped separately by Z and W chromosomes. While for the gene *IPO11*, whose Z/W recombination was suppressed after elegant-crested tinamou diverged from its sister species in Eurdomia S2 region, its gametologs are grouped by species (**Figure 4A**).

**Figure 4.**
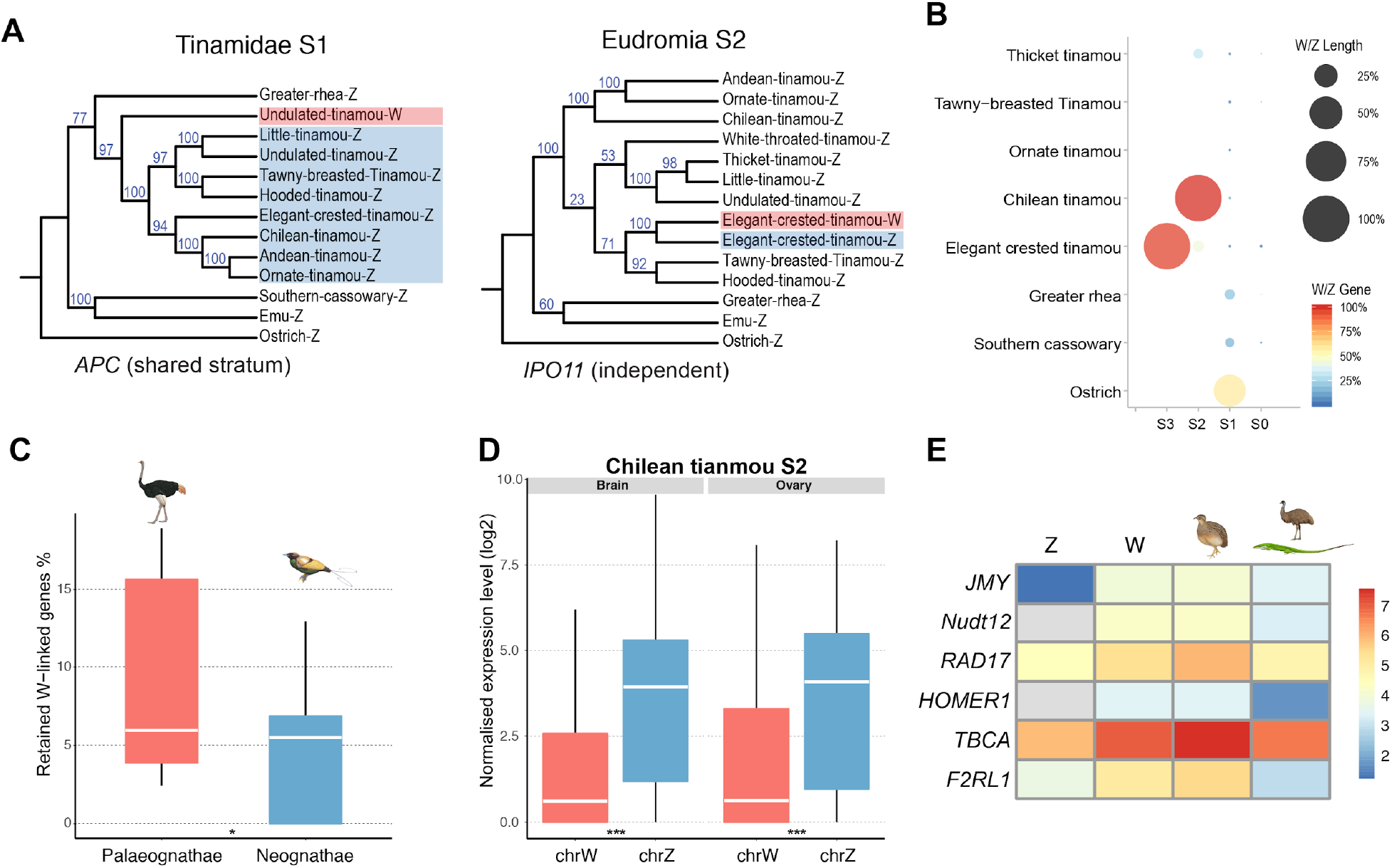
Evolution of sex-linked genes in Paleognathae. (A) Examples of gene trees for Z- and W-linked gametologs at different evolutionary strata. If sequences were grouped by their chromosomal origins, their residing evolutionary stratum was inferred to emerge before the species divergence. If sequences were grouped by their species origins, the stratum was inferred to emerge after the species divergence. We also labelled the chromosomal positions of the example genes at Figure 3A. (B) Comparison of W-linked genes and sequence lengths between different evolutionary strata. The areas of the circles are scaled to the W-to-Z ratios of the assembled lengths, and the colors of the circles are scaled to the W-to-Z ratios of the annotated genes. (C) Comparison of the percentages of retained W-linked genes at the SDRs of Palaeognathae vs. those of Neognathae. We show levels of significance (Wilcoxon test) with asterisks. *: 0.001=< *P*-value <0.05; (D) Comparison of gene expression levels between Z-linked and W-linked gametologs at the Chilean tinamou S2 evolutionary stratum. ***: *P*<0.0001. (E) The heatmap shows the normalized expression levels of Z-linked, W-linked gametologs, and also those of the entire genes of Chilean tinamou, as well as the mean expression levels of their orthologs in emu and green anole lizard.

Older evolutionary strata have lost more genes due to longer time of being suppressed for recombination (**Supplementary Table S9**). Both the assembled W-linked lengths, and the numbers of annotated W-linked genes relative to those of their Z-linked homologs dramatically decrease in the older evolutionary strata (**Figure 4B**). In general, paleognaths not only retained longer PARs (**Figure 3A**), but also retained a significantly (*P*<0.05, Wilcoxon test) higher percentage of W-linked genes at SDRs, relative to neognaths (**Figure 4C**). This indicated that the Paleognathae sex chromosomes not only underwent less frequent recombination suppressions, but also diverged from each other at a lower rate after the recombination was suppressed. Retained W-linked genes at SDRs show a decreased expression level relative to their Z-linked homologs among all the examined tissues in emu and Chilean tinamou with transcriptomes of both sexes available (**Supplementary Fig. S14**). This is consistent with the patterns of W-linked genes reported previously in other birds ^16,45^, or those of Y-linked genes in mammals ^20,46^, as evidence for functional degeneration after the RS. In particular, results from the extremely young Y chromosome of Drosophila ^47^ suggested that expression downregulation were likely to occur before the complete loss of genes on the W or Y chromosomes. This has also gained support in paleognathous birds by our close examination of genes in the youngest stratum of Chilean tinamou S2, where no W-linked genes have become deleted. Allele-specific expression analyses showed that over 50% of the genes have an at least 2-fold lower expression level on the W chromosome than on the Z chromosome (**Figure 4D, Supplementary Fig. S15**). Interestingly, we found that 6 of 119 expressed genes unexpectedly show a W-biased expression pattern in ovary, and also a higher overall expression level relative to their orthologs in the PAR or autosome of emu or lizard (**Figure 4E**). This indicated that these genes were specifically upregulated on the W chromosome. The orthologs of three genes, *JMY*, *Nudt2* and *TBCA* also have the highest expression level in ovary among all the human tissues^48^ (**Supplementary Fig. S16**). These results suggested that W-linked ovary-biased genes may undergo positive selection during their early stage of evolution, after RS restricted them in a female-specific environment.

The formation of evolutionary strata has also reshaped the TE landscape of the Z chromosomes, due to the reduction of average recombination rate from different time points, when the influenced part of Z chromosomes has stopped recombination in females. This has led to the differential accumulation of TEs on the Z chromosome. Overall, there is significantly higher TE content (*P*<0.01, Wilcoxon test) in the Z-linked SDR compared to PARs or autosomes, and also in the older strata compared to the younger ones (**Supplementary Fig. S17**). CR1 families that were active throughout the Palaeognathae evolution history are expectedly enriched on the Z chromosomes across all species, while some tinamou-specific families (e.g., CR1-J1/2_Pass) are only enriched on tinamou Z chromosomes (**Figure 2B**, **Supplementary Fig. S18**).

## Discussion

The ancestor of Palaeognathae was estimated to be probably volant and have a small body weighted between 2.9 to 5.5 kg ^6,15^, similar to that of extant tinamous. The convergent evolution of flight loss and gigantism ^1,49^ during Palaeognathae speciation have profound impacts on their genome evolution. The consequential deceleration of genome-wide substitution rates (**Figure 1A**) in the large-bodied ratites relative to tinamous probably has contributed to the different TE content (**Figure 2**) between ratites and tinamous, and also the much slower rate of divergence between sex chromosomes of ratites relative to those of any other birds. However, we demonstrated here that tinamous, with a similarly small body size and long branch length comparable to those of chicken, also exhibit long lengths of PARs and large presumably euchromatic W chromosome regions (**Figure 3A**). This is consistent with the previous cytogenetic studies, and a recent genomic study ^21^ that reported several tinamou species have homomorphic sex chromosomes^30^. It also indicated that substitution rates alone cannot explain the unique composition of Palaeognathae sex chromosomes.

We reasoned that the unique parental care mode of paleognathous species that originated in their ancestors probably account for their sex chromosome patterns. Palaeognathae have predominant (e.g. in ostrich) or exclusive (in tinamous) male-only parental care behaviors (**Supplementary Table S2**), which has been reported in about 1% of all extant bird species ^23,50^. Despite such rarity, paternal care, or the predominant bi-parental care (up to over 80%) of all extant bird species has been hypothesized to be the ancestral mode of parental care of birds ^23,50–52^. This was implicated by the study in shorebirds ^53^, and also in fish ^54^ showing that the transitions from paternal care to bi-parental or maternal cares are much more common than transitions from maternal care. It was also supported by the striking fossil discovery that paternal care probably also existed in the theropod dinosaurs closely related to birds ^24^. Paternal care is usually associated with reduced sexual selection targeting males, manifested as reduced levels of male-biased sexual size dimorphism and extrapair paternity ^26^. Indeed, the reversal of sex roles in parental care seems to result in a reversed pattern of sexual size dimorphism in most Paleognathous species with larger females than males (**Supplementary Table S2**). And unlike most other vertebrate species, it has been reported in emu that females would compete vigorously for access to males during mating season^55^. Although exclusive paternal care may seemingly restrict males’ opportunities for re-mating, behavioral and genetic studies showed that most paleognathous species are nevertheless polygynous or promiscuous ^50,56,57^. These results led us to propose that the evolution of paternal care in Palaeognathae probably help to ‘resolve’ the post-mating sexual conflict over care ^58^, and reduce the intensity of premating sexual selection, eventually produce the unique pattern of homomorphic sex chromosome pattern of Palaeognathae. Although slowly-diverging sex chromosomes have also been reported in several Neognathae species (e.g., killdeer and barn owl)^16^, they all have a bi-parental care system. More insights can be gained in future regarding the relationship between the parental care mode and sex chromosome evolution, by comparing species, for example, shorebirds that have recently evolved diverse modes of parental care ^59^.

It has been proposed that the aggregation of sexual antagonistic (SA) mutations nearby the sex-determining genes would select for the further expansion of RS between sex chromosomes ^27,28^, usually through chromosome inversions. Indeed, clear evidence of sequence inversions have been found for the most recent evolutionary stratum between the human X and Y chromosomes ^42,43^. However, a recent study in guppy fish pointed out that low levels of recombination is likely the cause rather than the result of the accumulation of SA mutations in this species ^60^. And studies in fungi species *Microbotryum lychnidis-dioicae* ^61^ and *Neurospora tetrasperma* ^62^ indicated that patterns of evolutionary strata can form without sexual antagonism or chromosome inversions. We have previously reported that the youngest evolutionary stratum of songbirds has probably formed due to the lineage-specific burst of TEs ^63^. In this work, we provided clear evidence that the youngest evolutionary stratum of Chilean tinamou formed without chromosome inversions between Z and W chromosomes (**Figure 3**). In this stratum, the majority of genes have down-regulated their expression levels in their W-linked gametologs relative to those of Z-linked gametologs (**Figure 4**). This occurred before the complete loss of substantial numbers of W-linked gametologs, and suggested that functional degeneration in the regulatory regions probably preceded that in the protein-coding regions after recombination was suppressed on the W or Y chromosomes. Some genes unexpectedly upregulated their expression levels in the ovary, i.e., underwent a ‘feminization’ process on the W chromosome. Both processes including the regulatory functional degeneration and sexualization (e.g., upregulation of Y-linked genes in testes) have previously been reported in the young Y chromosome systems of Drosophila species ^47,64,65^. These results together indicated that both are general characteristics of early sex chromosome evolution; and suggested that adaptive evolution of some genes with pre-existing sex-related functions, in response to the newly acquired sex-specific inheritance, may accelerate the degeneration of sex chromosomes by selective sweep ^66^.

## Materials and Methods

### Genome assembly and annotation

All animal procedures were carried out with the approval of China National Genebank animal ethics committee. We extracted genomic DNAs from the tissues of female samples of the studied species (**Supplementary Table S10**) using Gentra Puregene Tissue Kit (QIAGEN). Illumina paired-end DNA libraries were constructed and sequenced (Hiseq 4000) following the manufacturer’s protocol with varying insert sizes (250bp, 800bp, 2 or 5kb) for each species. We removed the reads containing more than 10% ambiguous nucleotides, or with more than 40% low-quality (Phred score <= 7) bases, or derived from adapter sequences and PCR duplications. We further trimmed the low-quality ends of reads if abnormal base compositions or low read qualities were detected. We then used the cleaned reads for *de novo* genome assembly using SOAPdenovo2^67^ (Version 2.04) and ALLPATHS-LG^68^ (r52488, with the parameter HAPLOIDIFY=True, a ploidy value set to two). Different k-mers were tested for SOAPdenovo2 and then the assembly results were compared to those of ALLPATHS-LG for each species to achieve the largest contig/scaffold N50 lengths. We then polished the assembly by Gapcloser (v1.12). We used Tandem Repeats Finder^69^ (TRF Version 4.09), RepeatProteinMask and RepeatMasker (version 4.0.6) with RepBase^70^ (20160829), and the *de novo* prediction program RepeatModeler (version open-1.0.8) together to predict and categorize the repetitive elements. For gene annotation, we first defined the candidate gene regions after aligning the query protein sequences of zebra finch, chicken and human (Ensembl release-87) to the targeted genomes using TBLASTN (E-value <= 1E-5). Then we refined the candidate gene regions by GeneWise^71^. 1000 such homology-based genes with a Genewise score of 100 were randomly selected to train Augustus^72^ (version 3.2.1) for *de novo* prediction. And we combined gene models from these two sources into a non-redundant gene set. The resulting gene models (measured by gene length, mRNA length, exon number and exon length) are comparable to those of other vertebrates. Gene names were further translated to those of their orthologs in the query species, based on the reciprocally best BLAST hit.

We first annotated the genomic copies of AviRTE for each of the Palaeognathae genome assembly using RepeatMasker (open-4.0.6) based on its published consensus sequences^35^. We then used all of the predicted AviRTE sequences from tinamou genomes to reconstruct the putative ancestral consensus sequence by Repeatmodeler (open-1.0.8), to calculate the divergence of AviRTE from each species. To date the time of horizontal transfer, we compared the flanking sequences (a block of Ns with same length) of each AviRTE copy between species by blat (version 36)^73^, using those of AviRTEs from *Crypturellus soui* with the highest number of copies as queries. We defined a pair of orthologous AviRTEs if their flanking sequences were aligned to each other.

We performed the PSMC^38^ analysis with the heterozygous SNP loci produced by GATK^74^. We set the parameters for the PSMC to be “N30 –t5 –r5 –p 4+30∗2+4+6+10,” and performed bootstraps (100 times) for each species to determine variance in the Ne estimates. We used the estimated values of generation time and mutation rate to scale the results. We used branch-specific estimates of the synonymous substitution rate per synonymous site (dS) from our dated phylogeny as proxies for rate of mutation. We used the age of sexual maturity collected from the published results, and multiplied by a factor of two as a proxy for generation time ^50^ (**Supplementary Table S2**).

### Phylogenomic analyses

The protein-coding genes from 3 amniotic species (*Gallus gallus*, *Anolis carolinensis* and *Homo sapiens* were downloaded from Ensembl (release-87). The gene sets from brown kiwi (PRJEB6383), white-throated tinamou and common ostrich (PRJNA212876) were obtained from NCBI. For gene loci with alternative splicing isoforms, only the longest transcript was retained. Then we used Treefam ^75^ to cluster genes from different species into gene families. 1). We performed all-versus-all alignments using BLASTP with E-value < 1E-7, and joined the fragmental alignments using Solar (a program in Treefam). The alignments were used to calculate the distance between two genes. Next, a hierarchical clustering algorithm was used to cluster all the genes, with the following parameters: min_weight=10, min_density=0.34, and max_size=500. 2). Multiple sequence alignment for each gene family was performed by MUSCLE ^76^ (v3.8.1551) and four-fold degenerate sites were extracted and concatenated to generate super-alignments. Our final dataset contained a total of 54,013 fourfold degenerate sites. We used MrBayes (v3.1.2) ^77^ to reconstruct the phylogeny with the (GTR + gamma) model using the following parameters: lset nst=6 rates=gamma; mcmc ngen=100000 printfreq=100 samplefreq=100 nchains=4 savebrlens=yes. For the NC tree, we first generated the whole genome alignments with LASTZ(1.04.00), using the genome of ostrich as the reference. In detail, each species’ genome was first built into pseudo-chromosome sequences using ostrich as reference, using a LASTZ parameter set designed for distant species comparison (--step=19 --hspthresh=2200 --inner=2000 --ydrop=3400 --gappedthresh=10000 --seed=12of19 for alignment, --minScore=1000 --linearGap=loose, with the scoring matrix HOXD55 for the chain step). The pairwise alignments were further processed by MULTIZ^78^ with the ROAST approach to construct multiple genome alignment, using a guided tree fixing the known phylogenetic positions of ostrich and chicken as outgroup. We then used the alignment filtering code (filter_alignment_maf_v1.1B.pl --window 36 --minidentity 0.55) described in ^9^ to remove the poorly-aligned regions. The multiple sequence alignments (MSAs) were then passed to MAFFT^79^ (--maxiterate 1000 --localpair) to obtain a further locally refined alignments. These steps resulted in 393,165,276 whole-genome aligned sites with 10.34 % gaps and undetermined sites, of which 15,565,579 distinct patterns can be used to estimate the phylogenetic topology. To construct the ML tree, we first preprocessed the non-informative sites with RAxML, the final MSA was first randomized into 100 subsets and each subset was used to generate a parsimony tree as the starting topology with RAxML^80^(8.2.11). Each starting tree was used together with its corresponding sequence subset to estimate a maximum likelihood tree with ExaML^81^ (3.0.19) under the GTRGAMMA model. Out of the 100 ML trees constructed above, the one with maximum ML was further used to evaluate the consistency and obtain the bootstrap value for the final tree. To further avoid the bias introduced by the protein-coding sequences, a genome-wide NC tree was further reconstructed based on the above whole genome alignments by excluding the coding sequences using msa_view in PHAST^82^ (v1.3). In total, 348,449,087 usable sites were obtained to construct the NC tree with the same approaches applied to the whole genome tree. The MCMCtree program (version 4.4) implemented in the Phylogenetic Analysis by Maximum Likelihood (PAML)^83^ package was used to estimate the species divergence time. Calibration time was obtained from the TimeTree database (http://www.timetree.org/). Two calibration points were applied in this study as normal priors to constrain the age of the nodes described below: 105-118 MA for the most recent common ancestor (TMRCA) of ostrich-chicken, and 271-286 MA for TMRCA of Aves and Dactyloidae. MitoZ^5^ was utilized with the default settings to assemble the whole mitochondrial genome for all studied Palaeognathae using a random subset (5 Gb) of paired-end reads from small insert-size library sequencing. The genes were identified using MITOS^84^ and curated by comparison with known sequences of other published ratites and tinamous from GenBank. Complete mitochondrial genomes were also used to construct a maximum likelihood phylogenetic tree using the same method as NC tree.

### Identification of sex-linked sequences

We identified the Z chromosome sequences out of the draft genomes based on their alignments to the Z chromosome sequences of ostrich, and also a female-specific reduction of read depth. Scaffold sequences of each species were aligned with LASTZ^85^ (version 1.02.00) to the ostrich Z chromosome sequence with parameter set ‘--step=19 --hspthresh=2200 --inner=2000 -- ydrop=3400 --gappedthresh=10000 --format=axt’ and a score matrix set for distant species comparison. Alignments were converted into a series of syntenic ‘chains’, ‘net’ and ‘maf’ results with different levels of alignment scores using UCSC Genome Browser’s utilities (http://genomewiki.ucsc.edu/index.php/). Based on the whole genome alignments, we first identified the best aligned scaffolds within the overlapping regions on the reference genome, according to their alignment scores with a cutoff of at least 50% of the whole scaffold length aligned in the LASTZ net results. We further estimated the overall identity and coverage distributions with a 10kb non-overlapped sliding window for each scaffold along the reference sequence, and obtained the distributions of identity and coverage. Scaffolds within the lower 5% region of each distribution were removed to avoid spurious alignments. Finally, scaffolds were ordered and oriented into pseudo-chromosome sequences according to their unique positions on the reference. Scaffolds were linked with 600 ‘N’s as a mark of separation. W-linked sequences are expected to also form an alignment with the reference Z chromosome of ostrich, but with lower numbers of aligned sequences and lower levels of sequence identity than their homologous Z-linked sequences, due to the accumulation of deleterious mutations after recombination was suppressed on the W. We also expect that there are still certain degrees (at least 70% as a cutoff) of sequence similarities between the Z- and W-linked sequences, for discriminating the true W-linked sequences from spurious alignments. After excluding the Z-linked sequences from the draft genome, we performed a second round of LASTZ alignment against the Z chromosome sequences of each species built from the above step. Then we excluded the spurious alignments with the cutoff of the pairwise sequence identity to be higher than 70%, but lower than 95%, and with the aligned sequences spanning at least 50% of the scaffold length. We further verified the W-linked sequences in species with sequencing data of both sexes available (**Supplementary Fig. S19**). We aligned the protein sequences of Z-linked genes with BLAT to the W-linked sequences, and then annotated the W-linked genes with GeneWise^71^. To construct the gametolog trees, CDS sequences of single-copy genes’ Z/W gametologs were aligned by MUSCLE^76^ and the resulting alignments were cleaned for ambiguous alignments by gblocks (-b4=5, -t=c, -e=-gb). Only the alignments longer than 300bp were used for constructing maximum likelihood trees by RAxML^80^ to infer whether their residing evolutionary stratum is shared among species or specific to certain lineages.

### Characterization of PARs and evolutionary strata

We aligned the raw reads of each species to their pseudo-chromosome sequences by BWA^86^ (0.7.12-r1039) with BWA-MEM algorithm. After removing PCR duplicates, we calculated read depth using SAMtools (version: 1.3.1) within each 100kb non-overlapping window and normalized it against the median value of depths per single base pair throughout the entire genome, to allow comparison among species. Boundaries of PAR were determined by a significant shift of depth values between neighboring windows. Similarly, we then scanned the Z/W pairwise alignment along the Z chromosome with a non-overlapping 100kb window to determine the boundaries between the neighboring strata, where there expected to be a sharp shift of the pairwise sequence identities or the occurrences of W-linked scaffolds. To determine the heterozygosity level of the evolutionary strata, we used the GATK^74^ (version 3.6-0-g89b7209) for SNP calling in both sexes for elegant crested tinamou and Chilean tinamou. SNPs whose read depths were too low (<100) or qualities lower than 100 were excluded. Boundaries of the very young stratum were determined by a significant shift of heterozygosity ratio between female and male. We used the method of assuming the molecular clock to infer the age of evolutionary strata, as well as relying on the species divergence time. In brief, we calculated branch-specific synonymous substitution rates (dS) of genes on the macrochromosomes by PAML (free-ratio model) for each species. The species-specific mutation rate to that of chicken (2.5×10-9/site/year) was scaled by their ratio of median values of the branch-specific synonymous substitution rates. We calculated the degree of male-biased mutation (α) by using the equation α=3(Z/A −2)/(4− 3Z/A), where ‘Z’ and ‘A’ represent the median substitution rates of intronic regions linked to autosomes and Z chromosome. The mutation rate of W chromosome is then estimated from the equation α=2A/W-1^87^, where ‘W’ refers to the mutation rate of the W chromosome. The sum of Z and W chromosome mutation rates in different strata results in the local divergence rate between the sex chromosomes. For the Z/W divergence level, the coding regions were removed from the pairwise Z/W genome alignments generated previously from LASTZ by customized perl scripts, and then we estimated the divergence level within non-coding regions by baseml in the PAML^83^ package after 1000 bootstraps. The age of a stratum was finally estimated by dividing the divergence level by divergence rate. Alternatively, the age of the strata was also estimated by the species divergence time, when we can map the strata to certain phylogenetic nodes.

We downloaded the transcriptomes of green anole lizard (brain, gonad: PRJNA381064) and Chilean tinamou (brain and gonad: PRJNA433114) from SRA. In addition, we collected the transcriptomes of adult emu brain, kidney, and gonad from both sexes. We used HISAT2 ^88^(version 2.1.0) for aligning the RNA-seq reads against reference genomes. Gene expression was measured by reads per kilobase of gene per million mapped reads (RPKM). To minimize the influence of different samples, RPKMs were adjusted by a scaling method based on TMM (trimmed mean of M values; M values mean the log expression ratios)^89^, which assumes that the majority of genes have similar expression levels across all samples. For the female transcriptomes from Chilean tinamou, we assigned the individual reads mapped to Chilean tinamou S2 region as W-linked or Z-linked based on the female-specific genomic SNP information. We used the ratios of RNA-seq read number vs. DNA-seq read number within a gene to normalize the mapping bias. W-linked or Z-linked spanning diagnostic SNPs within a gene were summed together to measure the allelic expression level.

## Acknowledgement

We would like to thank Gary Graves from Smithsonian Institute, Robb T. Brumfield and Donna L. Dittman from Louisiana State University Museum of Natural Science, Jack Withrow and Andy Kratter from Florida Museum of Natural History, Mariel L. Campbell and Ariel M. Gaffney from the Museum of Southwestern Biology, University of New Mexico for providing bird DNA samples for this work. Q.Z. is supported by National Natural Science Foundation of China (grant nos. 31722050, 31671319, 31861123001), the Fundamental Research Funds for the Central Universities (grant no. 2018XZZX002-04) and start-up funds from Zhejiang University. G.Z. is supported by Strategic Priority Research Program of the Chinese Academy of Sciences (XDB31020000, XDB13000000), International Partnership Program of Chinese Academy of Sciences (No. 152453KYSB20170002), Carlsberg foundation (CF16-0663), and Villum Foundation (No. 25900).

## Data availability

All the genomic reads, the genome assemblies and annotations generated in this study have been deposited in NCBI SRA and GenBank under the project accession number PRJNA545868. The above data have also been deposited in the CNGBdb (https://db.cngb.org/cnsa/) with accession number CNP0000505.

## References

1. Harshman, J. et al. Phylogenomic evidence for multiple losses of flight in ratite birds. Proc. Natl. Acad. Sci. U. S. A. 105, 13462–13467 (2008).

2. Cracraft, J. Phylogeny and evolution of the ratite birds. Ibis 116, 494–521 (1974).

3. Houde, P. Ostrich ancestors found in the Northern Hemisphere suggest new hypothesis of ratite origins. Nature 324, 563–565 (1986).

4. Cracraft, J. Continental drift, paleoclimatology, and the evolution and biogeography of birds. Journal of Zoology 169, 455–543 (1973).

5. Mitchell, K.J. et al. Ancient DNA reveals elephant birds and kiwi are sister taxa and clarifies ratite bird evolution. Science 344, 898–900 (2014).

6. Yonezawa, T. et al. Phylogenomics and morphology of extinct Paleognaths reveal the origin and evolution of the ratites. Curr. Biol. 27, 68–77 (2017).

7. Cloutier, A. et al. Whole-genome analyses resolve the phylogeny of flightless birds (Palaeognathae) in the presence of an empirical anomaly zone. Syst. Biol. (2019).

8. Smith, J.V., Braun, E.L. & Kimball, R.T. Ratite nonmonophyly: independent evidence from 40 novel Loci. Syst. Biol. 62, 35–49 (2013).

9. Jarvis, E.D. et al. Whole-genome analyses resolve early branches in the tree of life of modern birds. Science 346, 1320–1331 (2014).

10. Kapusta, A., Suh, A. & Feschotte, C. Dynamics of genome size evolution in birds and mammals. Proc. Natl. Acad. Sci. U. S. A. 114, E1460–E1469 (2017).

11. O’Connor, R.E. et al. Chromosome-level assembly reveals extensive rearrangement in saker falcon and budgerigar, but not ostrich, genomes. Genome Biol 19(2018).

12. Bromham, L. The genome as a life-history character: why rate of molecular evolution varies between mammal species. Philos. Trans. R. Soc. Lond. B Biol. Sci. 366, 2503–2513 (2011).

13. Mank, J.E. & Ellegren, H. Parallel divergence and degradation of the avian W sex chromosome. Trends Ecol. Evol. 22, 389–391 (2007).

14. Tollis, M. et al. Return to the sea, get huge, beat cancer: an analysis of cetacean genomes including an assembly for the humpback whale (Megaptera novaeangliae). Mol. Biol. Evol. 36, 1746–1763 (2019).

15. Berv, J.S. & Field, D.J. Genomic Signature of an Avian Lilliput Effect across the K-Pg Extinction. Syst. Biol. 67, 1–13 (2018).

16. Zhou, Q. et al. Complex evolutionary trajectories of sex chromosomes across bird taxa. Science 346, 1246338 (2014).

17. Takagi, N., Ito, M. & Sasaki, M. Chromosome studies in four species of Ratitae (Aves). Chromosoma 36, 281–291 (1972).

18. Lahn, B.T. & Page, D.C. Four evolutionary strata on the human X chromosome. Science 286, 964–967 (1999).

19. Nanda, I., Schlegelmilch, K., Haaf, T., Schartl, M. & Schmid, M. Synteny conservation of the Z chromosome in 14 avian species (11 families) supports a role for Z dosage in avian sex determination. Cytogenet. Genome Res. 122, 150–156 (2008).

20. Cortez, D. et al. Origins and functional evolution of Y chromosomes across mammals. Nature 508, 488–493 (2014).

21. Xu, L., Wa Sin, S.Y., Grayson, P., Edwards, S.V. & Sackton, T.B. Evolutionary dynamics of sex chromosomes of paleognathous birds. Genome Biol. Evol. (2019).

22. Li, W. Male-driven evolution. Curr. Opin. Genet. Dev. 12, 650–656 (2002).

23. Cockburn, A. Prevalence of different modes of parental care in birds. Proc. Biol. Sci. 273, 1375–1383 (2006).

24. Varricchio, D.J. et al. Avian paternal care had dinosaur origin. Science 322, 1826–1828 (2008).

25. Rice, W.R. Sex chromosomes and the evolution of sexual dimorphism. Evolution 38, 735 (1984).

26. Remeš, V., Freckleton, R.P., Tökölyi, J., Liker, A. & Székely, T. The evolution of parental cooperation in birds. Proc. Natl. Acad. Sci. U. S. A. 112, 13603–13608 (2015).

27. Rice, W.R. The accumulation of sexually antagonistic genes as a selective agent promoting the evolution of reduced recombination between primitive sex chromosomes. Evolution 41, 911–914 (1987).

28. Ponnikas, S., Sigeman, H., Abbott, J.K. & Hansson, B. Why do sex chromosomes stop recombining? Trends Genet. 34, 492–503 (2018).

29. Prum, R.O. et al. A comprehensive phylogeny of birds (Aves) using targeted next-generation DNA sequencing. Nature 526, 569–573 (2015).

30. Pigozzi, M.I. Diverse stages of sex-chromosome differentiation in tinamid birds: evidence from crossover analysis in Eudromia elegans and Crypturellus tataupa. Genetica 139, 771–777 (2011).

31. Reddy, S. et al. Why do phylogenomic data sets yield conflicting trees? Data type influences the avian tree of life more than taxon sampling. Syst. Biol. 66, 857–879 (2017).

32. Kimball, R.T. et al. A phylogenomic supertree of birds. Diversity 11, 109 (2019).

33. Hughes, A.L. & Hughes, M.K. Small genomes for better flyers. Nature 377, 391 (1995).

34. Suh, A. et al. Mesozoic retroposons reveal parrots as the closest living relatives of passerine birds. Nat. Commun. 2, 443 (2011).

35. Suh, A. et al. Ancient horizontal transfers of retrotransposons between birds and ancestors of human pathogenic nematodes. Nat. Commun. 7, 11396 (2016).

36. The origins of genome architecture. Choice Reviews Online 45, 45–0862 (2007).

37. Lynch, M. The Origins of Genome Architecture, 494 (Sinauer Associates Incorporated, 2007).

38. Li, H. & Durbin, R. Inference of human population history from individual whole-genome sequences. Nature 475, 493–496 (2011).

39. Abascal, F. et al. Extreme genomic erosion after recurrent demographic bottlenecks in the highly endangered Iberian lynx. Genome Biol. 17, 251 (2016).

40. Nadachowska-Brzyska, K., Li, C., Smeds, L., Zhang, G. & Ellegren, H. Temporal Dynamics of Avian Populations during Pleistocene Revealed by Whole-Genome Sequences. Curr Biol 25, 1375–80 (2015).

41. Barrón, M.G., Fiston-Lavier, A.-S., Petrov, D.A. & González, J. Population genomics of transposable elements in Drosophila. Annu. Rev. Genet. 48, 561–581 (2014).

42. Lemaitre, C. et al. Footprints of inversions at present and past pseudoautosomal boundaries in human sex chromosomes. Genome Biol. Evol. 1, 56–66 (2009).

43. Ross, M.T. et al. The DNA sequence of the human X chromosome. Nature 434, 325–337 (2005).

44. Tsuda, Y., Nishida-Umehara, C., Ishijima, J., Yamada, K. & Matsuda, Y. Comparison of the Z and W sex chromosomal architectures in elegant crested tinamou (Eudromia elegans) and ostrich (Struthio camelus) and the process of sex chromosome differentiation in palaeognathous birds. Chromosoma 116, 159–173 (2007).

45. Bellott, D.W. et al. Avian W and mammalian Y chromosomes convergently retained dosage-sensitive regulators. Nat. Genet. 49, 387–394 (2017).

46. Bellott, D.W. et al. Mammalian Y chromosomes retain widely expressed dosage-sensitive regulators. Nature 508, 494–499 (2014).

47. Zhou, Q. & Bachtrog, D. Chromosome-Wide Gene Silencing Initiates Y Degeneration in Drosophila. Current Biology 22, 522–525 (2012).

48. Consortium, G.T. et al. Genetic effects on gene expression across human tissues. Nature 550, 204–213 (2017).

49. Sackton, T.B. et al. Convergent regulatory evolution and the origin of flightlessness in palaeognathous birds. Science 364, 74–78 (2019).

50. Handford, P. & Mares, M.A. The mating systems of ratites and tinamous: an evolutionary perspective. Biol. Linn. Soc. 25, 77–104 (1985).

51. Wesolowski, T. On the origin of parental care and the early evolution of male and female parental roles in birds. Amer. Natur. 143, 39–58 (1994).

52. Tullberg, B.S., Ah-King, M. & Temrin, H. Phylogenetic reconstruction of parental-care systems in the ancestors of birds. Philos. Trans. R. Soc. Lond. B Biol. Sci. 357, 251–257 (2002).

53. Székely, T. & Reynolds, J.D. Evolutionary transitions in parental care in shorebirds. Proc. R. Soc. Lond. B Biol. Sci. 262, 57–64 (1995).

54. Gittleman, J.L. The phylogeny of parental care in fishes. Animal Behaviour 29, 936–941 (1981).

55. Coddington, C.L. & Cockburn, A. The Mating System of Free-Living Emus. Aust. Jour. Zoo. 43, 365 (1995).

56. Giraldo-Deck, L.M., Habel, J.C., Meimberg, H. & Garitano-Zavala, Á. Genetic evidence for promiscuity in the Ornate TinamouNothoprocta ornata (Aves: Tinamiformes). Bio. Linn. Soc. 120, 604–611 (2017).

57. Brennan, P.L.R. Mixed paternity despite high male parental care in great tinamous and other Palaeognathes. Animal Behaviour 84, 693–699 (2012).

58. Székely, T. Sexual conflict between parents: offspring desertion and asymmetrical parental care. Cold Spring Harb. Perspect. Biol. 6, a017665 (2014).

59. Reynolds, J.D. & Székely, T. The evolution of parental care in shorebirds: life histories, ecology, and sexual selection. Beha. Ecol. 8, 126–134 (1997).

60. Bergero, R., Gardner, J., Bader, B., Yong, L. & Charlesworth, D. Exaggerated heterochiasmy in a fish with sex-linked male coloration polymorphisms. Proc. Natl. Acad. Sci. 116, 6924–6931 (2019).

61. Branco, S. et al. Evolutionary strata on young mating-type chromosomes despite the lack of sexual antagonism. Proc. Natl. Acad. Sci. 114, 7067–7072 (2017).

62. Sun, Y., Svedberg, J., Hiltunen, M., Corcoran, P. & Johannesson, H. Large-scale suppression of recombination predates genomic rearrangements in Neurospora tetrasperma. Nat. Commun. 8, 1140 (2017).

63. Xu, L. et al. Dynamic evolutionary history and gene content of sex chromosomes across diverse songbirds. Nat Ecol Evol 3, 834–844 (2019).

64. Zhou, Q., Ellison, C.E., Kaiser, V.B., Alekseyenko, A.A. & others. .The epigenome of evolving Drosophila neo-sex chromosomes: dosage compensation and heterochromatin formation. PLoS 11 (2013).

65. Zhou, Q. & Bachtrog, D. Sex-specific adaptation drives early sex chromosome evolution in Drosophila. Science 337, 341–345 (2012).

66. Charlesworth, B. & Charlesworth, D. The degeneration of Y chromosomes. Philos. Trans. R. Soc. Lond. B Biol. Sci. 355, 1563–1572 (2000).

67. Luo, R. et al. SOAPdenovo2: an empirically improved memory-efficient short-read de novo assembler. Gigascience 1, 18 (2012).

68. Gnerre, S. et al. High-quality draft assemblies of mammalian genomes from massively parallel sequence data. Proc. Natl. Acad. Sci. U. S. A. 108, 1513–1518 (2011).

69. Benson, G. Tandem repeats finder: a program to analyze DNA sequences. Nucleic Acids Res. 27, 573–580 (1999).

70. Jurka, J. Repbase update: a database and an electronic journal of repetitive elements. Trends Genet. 16, 418–420 (2000).

71. Birney, E., Clamp, M. & Durbin, R. GeneWise and Genomewise. Genome Res. 14, 988–995 (2004).

72. Stanke, M., Steinkamp, R., Waack, S. & Morgenstern, B. AUGUSTUS: a web server for gene finding in eukaryotes. Nucleic Acids Res. 32, W309–12 (2004).

73. Kent, W.J. BLAT—The BLAST-Like Alignment Tool. Genome Res. 12, 656–664 (2002).

74. DePristo, M.A. et al. A framework for variation discovery and genotyping using next-generation DNA sequencing data. Nat. Genet. 43, 491–498 (2011).

75. Li, H. et al. TreeFam: a curated database of phylogenetic trees of animal gene families. Nucleic Acids Res. 34, D572–80 (2006).

76. Edgar, R.C. MUSCLE: multiple sequence alignment with high accuracy and high throughput. Nucleic Acids Res. 32, 1792–1797 (2004).

77. Ronquist, F. & Huelsenbeck, J.P. MrBayes 3: Bayesian phylogenetic inference under mixed models. Bioinformatics 19, 1572–1574 (2003).

78. Blanchette, M. et al. Aligning multiple genomic sequences with the threaded blockset aligner. Genome Res. 14, 708–715 (2004).

79. Katoh, K. & Standley, D.M. MAFFT multiple sequence alignment software version 7: improvements in performance and usability. Mol. Biol. Evol. 30, 772–780 (2013).

80. Stamatakis, A. RAxML version 8: a tool for phylogenetic analysis and post-analysis of large phylogenies. Bioinformatics 30, 1312–1313 (2014).

81. Kozlov, A.M., Aberer, A.J. & Stamatakis, A. ExaML version 3: a tool for phylogenomic analyses on supercomputers. Bioinformatics 31, 2577–2579 (2015).

82. Hubisz, M.J., Pollard, K.S. & Siepel, A. PHAST and RPHAST: phylogenetic analysis with space/time models. Brief. Bioinform. 12, 41–51 (2011).

83. Yang, Z. PAML 4: phylogenetic analysis by maximum likelihood. Mol. Biol. Evol. 24, 1586–1591 (2007).

84. Bernt, M. et al. MITOS: improved de novo metazoan mitochondrial genome annotation. Mol. Phylogenet. Evol. 69, 313–319 (2013).

85. Harris, R.S. Improved pairwise alignmnet of genomic DNA. (2007).

86. Li, H. & Durbin, R. Fast and accurate short read alignment with Burrows–Wheeler transform. Bioinformatics 25, 1754–1760 (2009).

87. Miyata, T., Hayashida, H., Kuma, K., Mitsuyasu, K. & Yasunaga, T. Male-driven molecular evolution: a model and nucleotide sequence analysis. Cold Spring Harb. Symp. Quant. Biol. 52, 863–867 (1987).

88. Kim, D., Langmead, B. & Salzberg, S.L. HISAT: a fast spliced aligner with low memory requirements. Nat. Methods 12, 357–360 (2015).

89. Robinson, M.D. & Oshlack, A. A scaling normalization method for differential expression analysis of RNA-seq data. Genome Biol. 11, R25 (2010).

90. HBW Alive: Handbook of the Birds of the World Alive. Choice Reviews Online 52, 52–0838 (2014).

